# Functional diversity alters the effects of a pulse perturbation on the dynamics of tritrophic food webs

**DOI:** 10.1101/2021.03.22.436420

**Authors:** Ruben Ceulemans, Laurie Anne Wojcik, Ursula Gaedke

## Abstract

Biodiversity decline causes a loss of functional diversity, which threatens ecosystems through a dangerous feedback loop: this loss may hamper ecosystems’ ability to buffer environmental changes, leading to further biodiversity losses. In this context, the increasing frequency of climate and human-induced excessive loading of nutrients causes major problems in aquatic systems. Previous studies investigating how functional diversity influences the response of food webs to disturbances have mainly considered systems with at most two functionally diverse trophic levels. Here, we investigate the effects of a nutrient pulse on the resistance, resilience and elasticity of a tritrophic—and thus more realistic—plankton food web model depending on its functional diversity. We compare a non-adaptive food chain with no diversity to a highly diverse food web with three adaptive trophic levels. The species fitness differences are balanced through trade-offs between defense/growth rate for prey and selectivity/half-saturation constant for predators. We showed that the resistance, resilience and elasticity of tritrophic food webs decreased with larger perturbation sizes and depended on the state of the system when the perturbation occured. Importantly, we found that a more diverse food web was generally more resistant, resilient, and elastic. Particularly, functional diversity dampened the probability of a regime shift towards a non-desirable alternative state. In addition, despite the complex influence of the shape and type of the dynamical attractors, the basal-intermediate interaction determined the robustness against a nutrient pulse. This relationship was strongly influenced by the diversity present and the third trophic level. Overall, using a food web model of realistic complexity, this study confirms the destructive potential of the positive feedback loop between biodiversity loss and robustness, by uncovering mechanisms leading to a decrease in resistance, resilience and elasticity as functional diversity declines.

## 1 Introduction

Human activities undeniably disrupt ecosystem structure and functioning (Hooper et al., 2005; Worm et al., 2006; Cardinale et al., 2012; Hautier et al., 2015). Direct effects such as habitat loss due to pollution (Dudgeon et al., 2006; Butchart et al., 2010; Hölker et al., 2010) and increased land requirements for agricultural or industrial use (Brooks et al., 2002; Ryser et al., 2019; Horváth et al., 2019) are major causes of the observed losses in biodiversity worldwide. Moreover, climate change effects have a decisive influence on these losses (Bestion et al., 2020): in addition to the global temperature rise (Hansen et al., 2006), the frequency of disruptive extreme weather events has increased steadily (Easterling et al., 2000). For instance, recurrent storms or heavy rainfalls amplify excessive nutrient loading in rivers, lakes, and coastal areas, causing species losses (Øygarden et al., 2014). The combined effect of these processes on biodiversity creates a potentially dangerous feedback loop. When biodiversity is lost, the respective decrease in functional diversity may alter the ecosystems ability to buffer perturbations (Cardinale et al., 2012; García-Palacios et al., 2018; Ceulemans et al., 2019). This leads to additional biodiversity losses, and consequently to more vulnerable ecosystems.

In the last decades, the functional perspective has brought new insights to quantify the effects of biodiversity loss on ecosystem functioning (Violle et al., 2007; Tirok and Gaedke, 2010). One approach entails sorting out the members of a food web into functional groups with similar trait values or to consider explicitly the trait values and their distributions within trophic levels. In this way, morphological, physiological or behavioral individual characteristics are linked to a certain function, such as growth rate or nutrient uptake (Garnier, Navas, and Grigulis, 2016), and depend on each other by trade-offs to determine the overall fitness (McGill et al., 2006; Violle et al., 2007). This approach makes explicit how trait changes can feed back to population and food web dynamics, and partly regulate the response of food webs to environmental changes (Yamamichi and Miner, 2015; Theodosiou, Hiltunen, and Becks, 2019; Raatz, Velzen, and Gaedke, 2019).

Most studies investigating the responses of food webs to perturbations are restricted to multitrophic systems where only one or two trophic levels may adapt (Persson et al., 2001; Kovach-Orr and Fussmann, 2013), or to strictly bitrophic systems (Jones, 2008; Fussmann and Gonzalez, 2013; Yamamichi, Yoshida, and Sasaki, 2011; Bell et al., 2019; Govaert et al., 2019; Raatz, Velzen, and Gaedke, 2019). These studies underline how functional diversity at one or two trophic levels generally enhances the ecosystem’s ability to buffer against perturbations, and that the effects of functional diversity can be modulated at different trophic levels. However, tritrophic systems with at least two functionally diverse trophic levels are more realistic, since strictly bitrophic interactions are rare in nature (Pimm et al., 2014; Matsuno and Nobuaki, 1996; Abdala-Roberts et al., 2019), and top predators can have a large influence on ecosystem functioning and dynamics (Estes et al., 2011; Brose et al., 2019; Ceulemans, Guill, and Gaedke, 2020).

To properly understand how tritrophic food webs respond to environmental changes, insights are needed into the mechanisms driving their response. External disturbances come in a variety of forms, and each can affect the food web and its functions in different ways. Broadly, external disturbances can be separated into two types, called press and pulse perturbations (Tilman and Downing, 1996; Raatz, Velzen, and Gaedke, 2019). Press perturbations are long-term or permanent changes to a system component, such as increased harvesting or warming. In contrast, pulse perturbations are short-term and quasi-instantaneous changes to state variables, such as species biomasses or nutrient concentration, e.g. due to a forest fire or massive rainfall causing heavy run-off (Bender, Case, and Gilpin, 1984; Harris et al., 2018).

In this study, we investigate the effects of a nutrient pulse on the dynamics of tritrophic food webs with different amounts of biodiversity. Nutrient pulses correspond to a temporary boost of the carrying capacity, which can destabilize the dynamics of food webs and put species at increased risk of extinction (Rosenzweig, 1971). In aquatic systems, such events are happening with increased frequency and magnitude (Galloway et al., 2008; Kaushal et al., 2014). This is highly worrying, because eutrophication leads to drastic reduction in water quality (Kaushal et al., 2014; Couture et al., 2018; Díaz et al., 2019), and to the apparition of anoxic dead zones (Diaz and Rosenberg, 2008). Human-induced nutrient loading changes the usual seasonal pattern of phytoplankton and zooplankton (Sommer et al., 2012). Importantly, by affecting the timing and the amplitude of plankton biomass peaks, the dynamics of the upper trophic levels may be strongly affected as well (Cloern and Jassby, 2008). Moreover, sudden increases in available nutrients can lead to changes in ecosystem functions, which may be difficult to reverse (Carpenter, 2005). The probability of such a regime shift may be influenced by the amount of diversity present in the ecosystem (Folke et al., 2004; Ceulemans et al., 2019), but explicit demonstration of the mechanisms by which this happens remains difficult.

The response of a tritrophic food web to a nutrient pulse is characterized by several aspects, which are measured by different quantities. Analogous to Grimm and Wissel, 1997 and Raatz, Velzen, and Gaedke, 2019, the following terms are used:

- **Resistance** refers to the maximum temporary change in dynamics after a pulse perturbation.
- **Resilience** refers to whether or not the system returns to its original state after a pulse perturbation.
- **Elasticity** refers to how quickly the system returns to its original state.

These three quantities are evaluated by several properties of the food web dynamics. The resistance is evaluated shortly after the perturbation. When the dynamics are strongly affected before the system returns to its original state, the resistance is low. The resilience is simply determined by examining the dynamics after a long time period following the perturbation. If the system does not return to its original state, it is not resilient. Finally, the elasticity is estimated through the return time, which is the time it takes for the system to return to the original state. A lower return time corresponds to a higher elasticity.

We investigated the response of tritrophic food webs with low and high functional diversity to a temporal nutrient increase, by comparing a tritrophic non-adaptive food chain to a food web which is adaptive at each trophic level. We used a food web model, which is adaptive in the sense of species sorting, as described in Ceulemans et al. (2019). Prey species are either defended or undefended, and predator species are either selective or non-selective feeders (Fig. 1). Their relative importance changes according to ambient conditions, leading to continual changes of the mean trait values at each trophic level. Our hypotheses are that the food web response depends on (i) the perturbation size, (ii) the time at which the perturbation is applied, and (iii) on the functional diversity. The last hypothesis implies that an adaptive food web is less affected by a nutrient pulse than a non-adaptive food chain. To enlarge generality and accurately capture the complex behavior of the system, we study this response in different parameter regions with multiple attractors by varying the Hill exponent of the functional responses.

**Figure 1.**
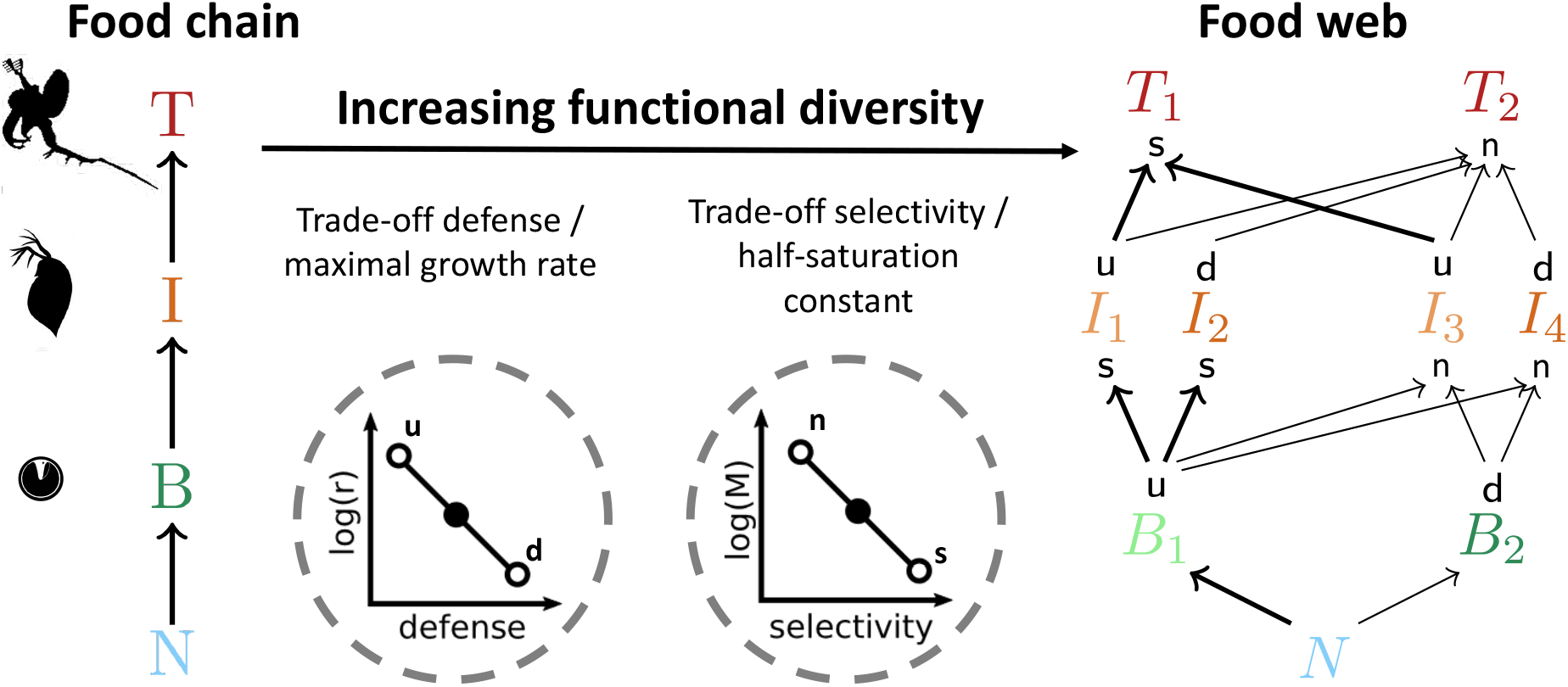
Comparison of two food webs with different functional diversity. The system with no functional diversity (left side, “chain”) is a non-adaptive food chain, where nutrients (*N*) are taken up by a basal species (*B*), which is consumed by an intermediate species (*I*), which is preyed on by a top species (*T*). In the highly diverse system (right side, “web”), prey species are either undefended (`u`), or defended (`d`), and predators are either selective (`s`) or non-selective (`n`) feeders. The basal and the top species each differ by one trait, whereas the intermediate species, being both consumers and prey, differ by two traits, resulting in four functionally unique species. Two trade-offs are used to balance the fitness of the species: a higher defense comes at the cost of a lower growth rate (*r*), and being less selective implies a larger prey spectrum, but also an increased prey uptake half-saturation constant (*M*). In this way, a defended species grows slower than an undefended one and a selective feeder can more efficiently exploit low prey concentrations. The resulting differences in trophic interaction strengths are shown by the arrow thickness between the species.

## 2 Methods

As a basis for our study, we used the chemostat tritrophic model described in Ceulemans et al., 2019, consisting of free inorganic nutrients (nitrogen, *N*) and three trophic levels: basal (*B*), intermediate (*I*) and top (*T*). In this model, the functional diversity of the food web can differ by changing the trait differences between the species on all trophic levels. In this study, we consider two distinct cases: a non-adaptive food chain with no functional diversity; and the highly diverse food web, where all the trophic levels are adaptive in the sense of species sorting (see Fig. 1). Species have fixed traits and adaptation occurs through the different responses of the asexually reproducing species. For convenience, we will call the food web with no functional diversity “chain” and the highly diverse food web “web” in the following. In the latter, the species’ fitness varies with the trait difference and arises from two trade-offs. Prey species are defended (*d*) or undefended (*u*) against predation, but defense comes at the cost of a slower growth rate (*r*). Predator species are selective (*s*) or non-selective (*n*). Selectivity implies preferential feeding on undefended species, and more efficient exploitation of low resource biomass densities due to a lower half-saturation constant (*M* for *I* and *µ* for *T*, cf. Eqs. (4)–(5)). The chain has three species (one per trophic level), and the web has nine species: two basal species, four intermediate species and two top species (Fig. 1). The intermediate trophic level has four species since they are both prey and predator species, therefore they have two distinct traits, which results in four possible combinations. In all equations, we use these equivalences:

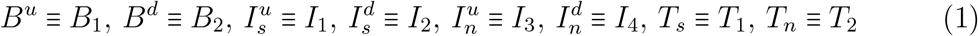

The tritrophic model described in Ceulemans et al., 2019 is written as followed:

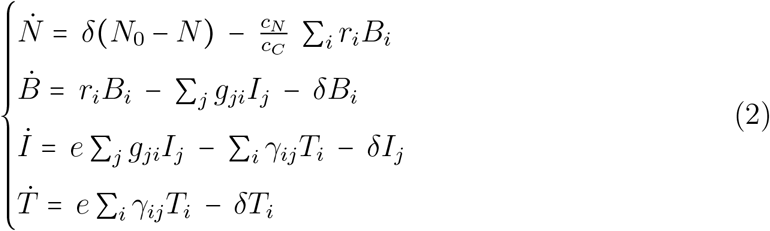

In these equations, the species are distinguished by *i* ∈ {1, 2} at the basal and top trophic level, and *j* ∈ {1, 2, 3, 4} at the intermediate trophic level. To increase realism, we assume a flow-through chemostat system rather than a static batch situation. Nitrogen is the limiting nutrient (*N*), with incoming concentration *N*_0_. The dilution rate is set by *δ*. Since the nutrients are measured in nitrogen concentration and the species biomass in carbon biomass, the nitrogen-to-carbon weight ratio 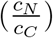 scales the basal (*B*_*i*_) growth terms in the nutrient equation. The following equations give the expressions of the basal growth rate *r*_*i*_ and of the basal-intermediate and intermediate-top functional responses *g*_*ji*_ and *γ*_*ij*_ respectively.

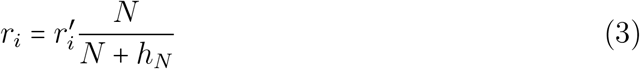

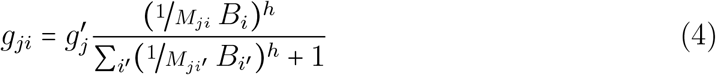

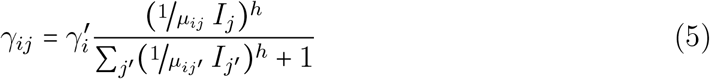

where *r′* denotes the maximal basal growth rate and *h*_*N*_ the nutrient uptake half-saturation constant; *g′* and *γ′* denote the maximal grazing rates of *I* and *T, M* and *µ* the half-saturation constants of the *B* − *I* and *I* − *T* interaction, and *h* the Hill exponent.

The two trade-offs affect the trait values in the above equations. For the prey, the basal species has a growth rate 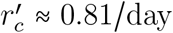 in the chain, whereas in the web, the undefended basal species’ growth rate 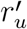 is set to 1/day, and for the defended species 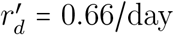. For the predators, the intermediate and top species in the chain can

exploit their prey with a half-saturation constant *M*_*c*_ = *µ*_*c*_ ≈ 424 *µgC* / *L*, respectively. For the web, the intermediate and top selective feeders have half-saturation constants set to *M*_*s*_ = *µ*_*s*_ = 300 *µgC*/*L* for the undefended prey, and the corresponding value for the non-selective feeders on all prey types is = *M*_*n*_ = *µ*_*n*_ = 600 *µgC / L* (See A1.1, Appendix for all the parameter values with units, which reflect a typical plankton food web, detailed legitimations are provided in Ceulemans et al., 2019).

Importantly, our previous study showed that the Hill exponents of the functional responses of the *B* − *I* and *I* − *T* interaction (*h*, cf. Equations (4)–(5)) play an important role in determining the nature of the dynamical attractor to which the system relaxes Ceulemans et al. (2019). In particular, two attractors exist for both the chain and the web. The “low production state” (LP) has a low top biomass production and large biomass oscillations, whereas the “high production state” (HP) has a high top biomass production and small biomass oscillations. In some cases the chain and the web exhibit bistability (see Table 1) and are potentially under the threat of a regime shift following a perturbation. In our model, we changed the Hill exponent to capture the different dynamical patterns and to consider all possible behaviours of the model after the perturbation. The Hill exponent is kept above 1.05 to make sure that all the species coexist. In particular we have selected three values of Hill exponents (*h* = 1.05, *h* = 1.10 and *h* = 1.15) for which we will investigate the system’s response.

**Table 1:** Summary of the effect of the three values of Hill exponent on the attractors found in the chain and web. The system either relaxes to the low-production state (*LP*), the high production state (*HP*), or is bistable. A visual representation of the dynamics on each attractor is given in Fig. A3.1, Appendix. In this text, we distinguish between attractor *type*, denoting whether the attractor is a fixed point, limit cycle, or chaotic, and attractor *shape*, distinguishing between the *HP* or *LP* state. By varying the Hill exponent, we can investigate the effect of a nutrient pulse perturbation under different dynamical regimes.

| Hill exponent | Attractor and structure |  |
| --- | --- | --- |
|  | chain | web |
| 1.05 | *LP*: limit cycle | *LP*: chaotic<br>*HP*: limit cycle |
| 1.10 | *LP*: limit cycle<br>*HP*: limit cycle | *HP*: limit cycle |
| 1.15 | *HP*: limit cycle | *HP*: fixed point |

To evaluate the system’s response to a nutrient pulse, we always ensured that the system is at an attractor (Fig. 2), and not still in a transient state. Importantly, the attractor may have a more complex structure than a simple fixed point: it can be a limit cycle, or a chaotic attractor. In the latter cases, the individual populations do not settle down to a fixed value but remain oscillating perpetually. When the system has relaxed to the attractor, the perturbation is applied at time *t*_*P*_ by altering the free nutrient concentration *N* :

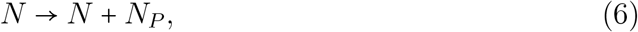

where *N*_*P*_ is the amount of added extra nutrients, also referred to as the perturbation size. This instantaneous change of the state variable *N* moves the system from its former location on the attractor to a point farther away from it (Fig. 2).

**Figure 2.**
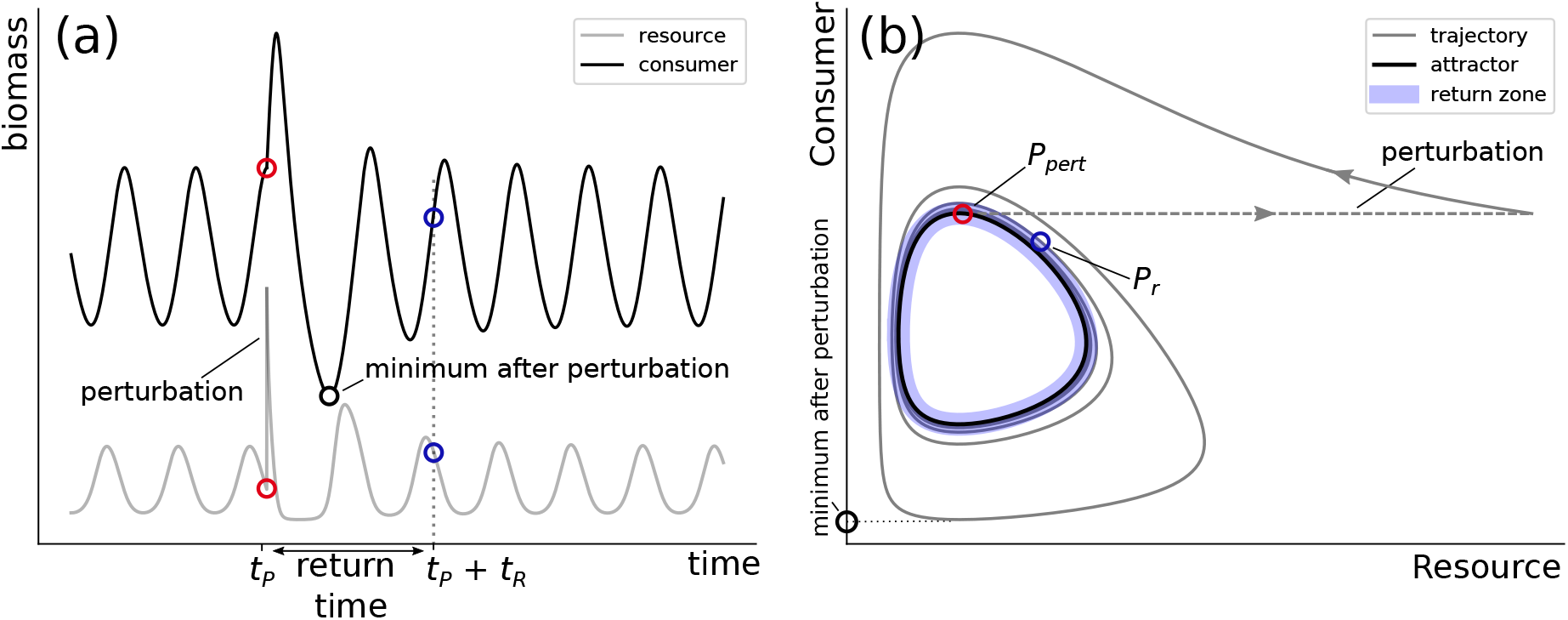
Schematic graphical examples of the different quantities calculated for our results. To study the ecosystem response to a perturbation, we recorded the minimal biomasses reached by the populations after the perturbation, as well as the time it took for the trajectory to return to the vicinity of the attractor. Panel (a) shows the timeseries of a simple consumer-resource model. The system is oscillating on its stable limit cycle until a nutrient pulse perturbation is applied at time *t*_*P*_ (red circle), after which the consumer increases and then declines to very low values. Panel (b) shows the same timeseries data but plotted in phase space. The limit cycle on which the system is oscillating originally is shown by the black curve. The nutrient pulse perturbation is applied at the red point *P*_pert_ (also shown in panel (a)), after which the system relaxes back to the attractor. The return time *t*_*R*_ is measured by the time it takes for the trajectory to remain inside the close neighborhood of the attractor (return zone, indicated by the colored region). This happens at the blue point *P*_*r*_ (also indicated in panel (a)).

After the perturbation, there are three possible outcomes for the perturbed system. The first option is that the pulse perturbation temporarily disturbs the system, by altering its dynamics during a transient phase in which the system relaxes back to the attractor. In this case, we recorded the time it took for the system to return to the attractor (return time *t*_*R*_), as well as the biomass minima and maxima of each population and each trophic level as a whole during this phase (see also Fig. 2). A second outcome may occur when at least one population biomass reaches such a low value that it crosses the extinction threshold set to 10 ^−9^ *µgC / L*. Below this value, the population is set to 0 and is considered extinct. Consequently, the system can never return to the initial attractor. Such a threshold prevents numerical problems that can occur when state variables reach values extremely close to 0. Finally, a third outcome is possible when the system is bistable. In that case, it may happen that the trajectory is pushed inside the other attractor’s basin of attraction. Therefore, the system also never returns to the initial system, but all populations are still present in the food web.

Additionally, when the attractor is more complex than a fixed point (i.e., a limit cycle or chaotic attractor), we investigated how the effect of the perturbation on the dynamics depends on where on the attractor it is applied. This means that for each point on the attractor, we perturbed the system and calculated the biomasses’ minima and maxima as well as the return time following this perturbation. For this, we sampled the different attractors in a high spatial resolution. This was achieved in multiple steps. First, starting from an initial condition known to be in the basin of attraction of the relevant attractor, the system was allowed to relax for 10^5^ time units using a large timestep Δ_*t*_ = 10 ^−1^ such that a point sufficiently close to the attractor could be obtained. If the attractor was a limit cycle, the system was further integrated for approximately one period using a high temporal resolution (Δ_*t*_ = 10^−3^). This creates a set of points 𝒮_*t*_ on the attractor. Finally, to sample the attractor in a way such that the distances between the sampled points on the attractor do not become very large when the dynamics are moving very fast, the attractor was interpolated and resampled such that the arc length between consecutive sampled points is equal to 1 (in units of biomass). For this set of spatially sampled points 𝒮_*x*_, the distance between a point on the attractor and the closest point to it in 𝒮_*x*_ is guaranteed to be smaller than 1 *µgC L*. In our results, we defined the return zone (cf. Fig. 2) as the volume in phase space inside which the distance to any point on the attractor is less than 5 *µgC L*). When the attractor is chaotic, even when performing this procedure over multiple periods, the return time could not accurately be calculated using this method. Hence, we excluded results presenting chaotic attractors in the elasticity analysis.

## 3 Results

We compared the response of a tritrophic non-diverse chain (“chain”) and a diverse food web (“web”) to a nutrient pulse, by quantifying its resistance, resilience, and elasticity (see Figs. 1-2). The major results are summarised schematically in Fig. 3. Alongside this analysis, we investigated the actual timeseries in detail in order to uncover the mechanisms responsible for the observed responses.

**Figure 3.**
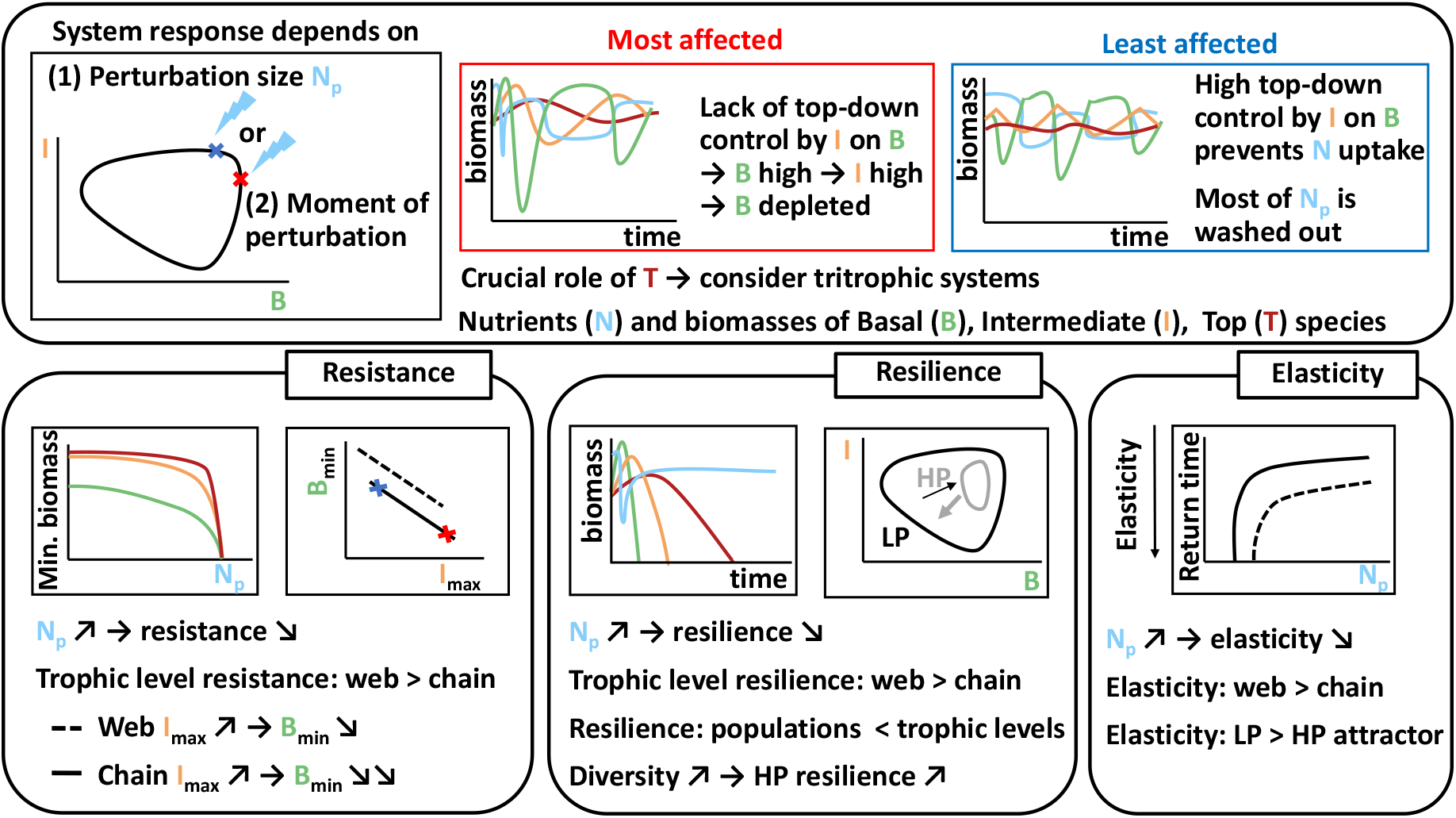
Sketch of the response of a tritrophic food web to a nutrient pulse. This response depends on the amount of extra nutrients added (the perturbation size *N*_*p*_), and on the moment at which the perturbation is applied (i.e., the position on the attractor). Two extreme areas on the attractor (i.e., two different specific moments of perturbation) could be distinguished: the most (red) and the least (blue) affected locations in phase space. The three studied characteristics—resistance, resilience and elasticity—show different aspects of how a food web responds to a nutrient pulse perturbation, depending on the Hill exponent, the nature of the attractor (cf. Table 1, *HP* : high-production attractor, *LP* : low-production attractor) and its functional diversity.

### 3.1 Resistance

As a general pattern, the resistance, here quantified by the biomass minima reached after a perturbation (cf. Fig. 2) tend to decrease as the perturbation size *N*_*P*_ (i.e the added extra nutrients) increases (Fig. 4). This implies that, as the amount of added nutrients increases, the biomass amplitudes immediately after the perturbation increase correspondingly, in a highly nonlinear way (see also Appendix Fig. A3.2 showing an increase in the maxima as well). We observe that, in all cases, the basal level is most strongly affected by the nutrient pulse, whereas the top level is least affected.

**Figure 4.**
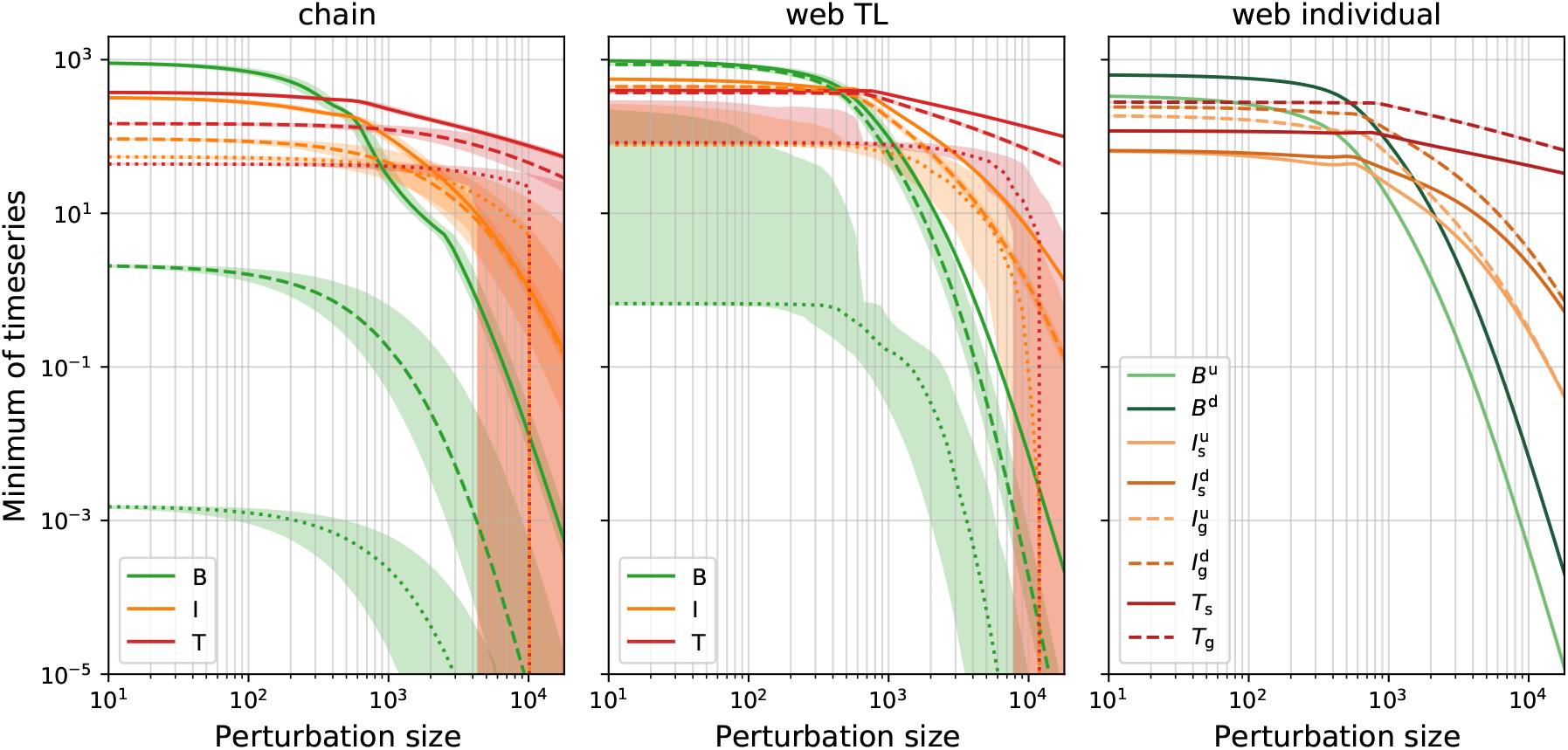
Minima reached by the timeseries after the perturbation, for the chain (left), the total biomass per trophic level in the diverse food web (middle), and the individual populations in the food web (right), as a function of the perturbation size. For the chain and trophic level biomass in the food web, the solid lines show the minima for Hill exponents *h* = 1.15, dashed for *h* = 1.10, and dotted for *h* = 1.05. The individual populations minima for the food web are only shown in the case of *h* = 1.05. Each line corresponds to the median of 1000 different randomly sampled initial conditions that lead to coexistence of all species, with the shaded area showing the upper and lower quartiles. These initial conditions were first allowed to relax to the attractor for 3 · 10^4^ time units before the perturbation was applied.

By randomly selecting 1000 time points at which the perturbation is applied for each perturbation size, we capture how the system response depends on the specific time—and thus the state of the system—at which the perturbation occurs. Importantly, this analysis shows how the shape of the attractor influences its resistance. If the equilibrium state is a fixed point (as is the case for the web at *h* = 1.15), the response does not depend on the moment of perturbation. All initial conditions lead exactly to the same post-perturbation minimum and therefore the median, upper, and lower quartiles are all equal (Figure 4). On the other hand, when the equilibrium state is a limit cycle (or a chaotic attractor), the minimum reached by the timeseries may strongly vary depending on where on the limit cycle the perturbation is applied. How much this response varies is reflected by the difference between the upper and lower quartiles.

When applying only a very small perturbation to the food web, the spread of the biomass minima is correspondingly small (Fig. 4). However, this spread appears very large for the web when *h* = 1.05 (Fig. 4, middle panel). This apparent discrepancy can be explained by the presence of the two attractors (cf. Table 1), each with a basin of attraction of approximately equal size. Because the actual timeseries minima differ between the attractors, so do the minima after a small perturbation. Thus, what appears as a very large range between which the minima are distributed, is actually a strongly bimodal distribution centered around the respective minima of each attractor. While the chain for *h* = 1.1 also exhibits bistability (with both the high and low production attractor), the basin of attraction of the high production attractor is so small that this effect does not significantly impact our results (only one initial condition out of 1000 ended up on the high production attractor).

The pronounced differences between the minimal biomass values reached after a perturbation of given size can be understood by explicitly examining the responses when the perturbation occurs at different states of the system (i.e along the attractor, Fig. 5 for *h* = 1.10) and the post-perturbation timeseries (Fig. 6). In these figures, we highlight six points on the attractors, and compare the response of the basal trophic level to a large perturbation of size 10^4^ *µgN* / *L* (≈ 10. *N*_0_ the standard nutrient inflow) applied at each of these points. While *P*_1_ and *P*_2_ are very close together on the attractor, the effect of the perturbation on the resulting dynamics is very different between these two points (Fig. 5, 6a–c). In both cases, the basal species are in decline at the moment of the perturbation. At *P*_1_ (Fig. 6b), they are under sufficient top-down control by the intermediate level, such that the free nutrients cannot be efficiently exploited and remain very high for a long period of time. Thus, most of the extra nutrients due to the perturbation are simply washed out of the system. In the case of *P*_2_ (Fig. 6c), however, the basal species are able to exploit almost all of the newly available nutrients immediately. This leads to an extremely high peak biomass of the basal level, which is in turn exploited by the intermediate level. Because of the delayed response of the top level, the intermediate species are able to stay at a high biomass for an extended period of time and thus graze the basal level down to a very low biomass density.

**Figure 5.**
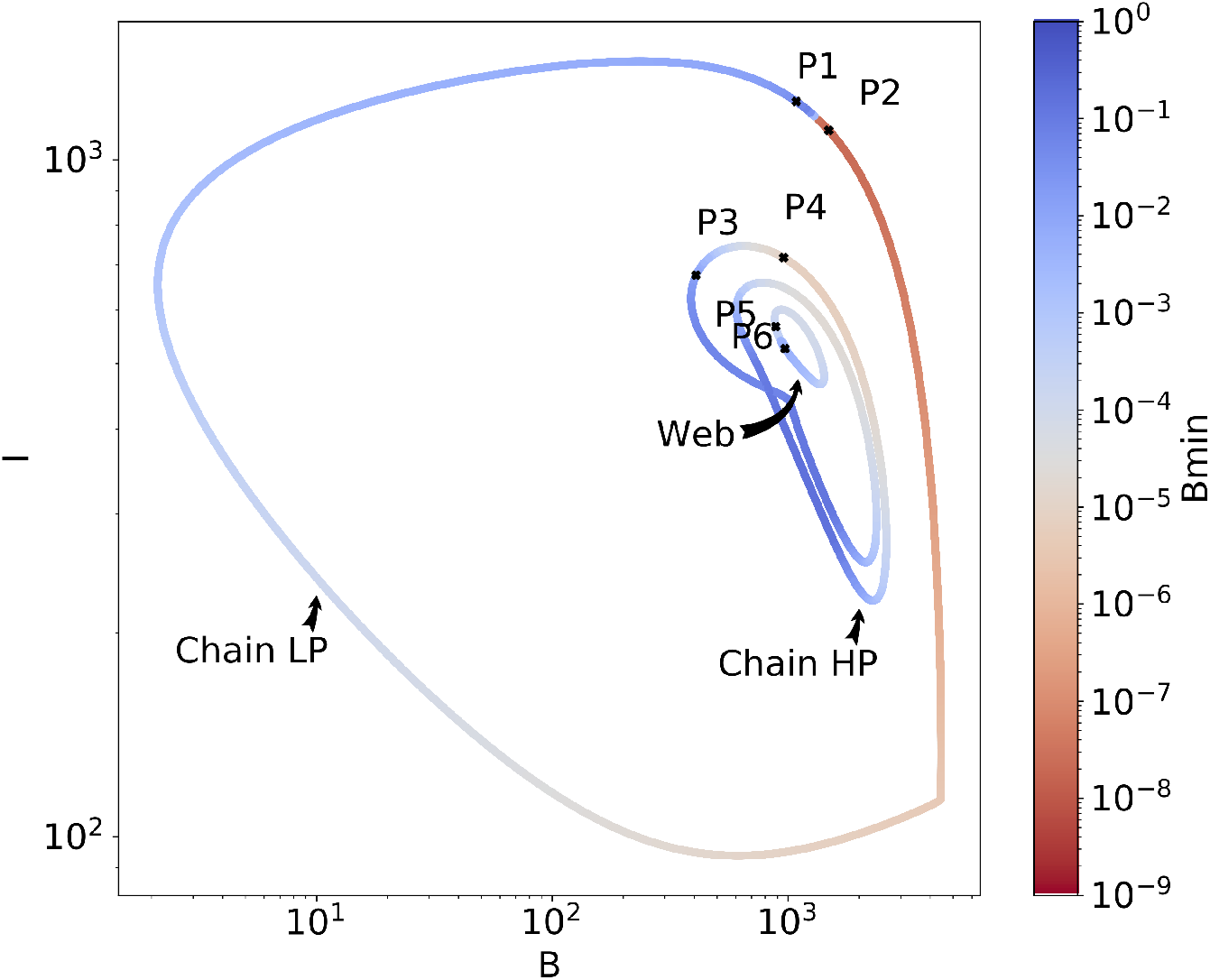
Minima reached by the basal trophic level after the perturbation, depending on where on the attractor a nutrient pulse of size 10^4^ is applied, projected on the *B* − *I* plane. The two attractors of the chain (*LP, HP*), and the *HP* attractor of the web, when *h* = 1.1, are all shown. The points *P*_1_ to *P*_6_ are picked to highlight the large differences in response that are possible by perturbing the system on points that may be very close together.The timeseries of these points are shown in Fig. 6. See Fig. A3.3, in the Appendix for all Hill exponents and for different perturbation sizes.

**Figure 6.**
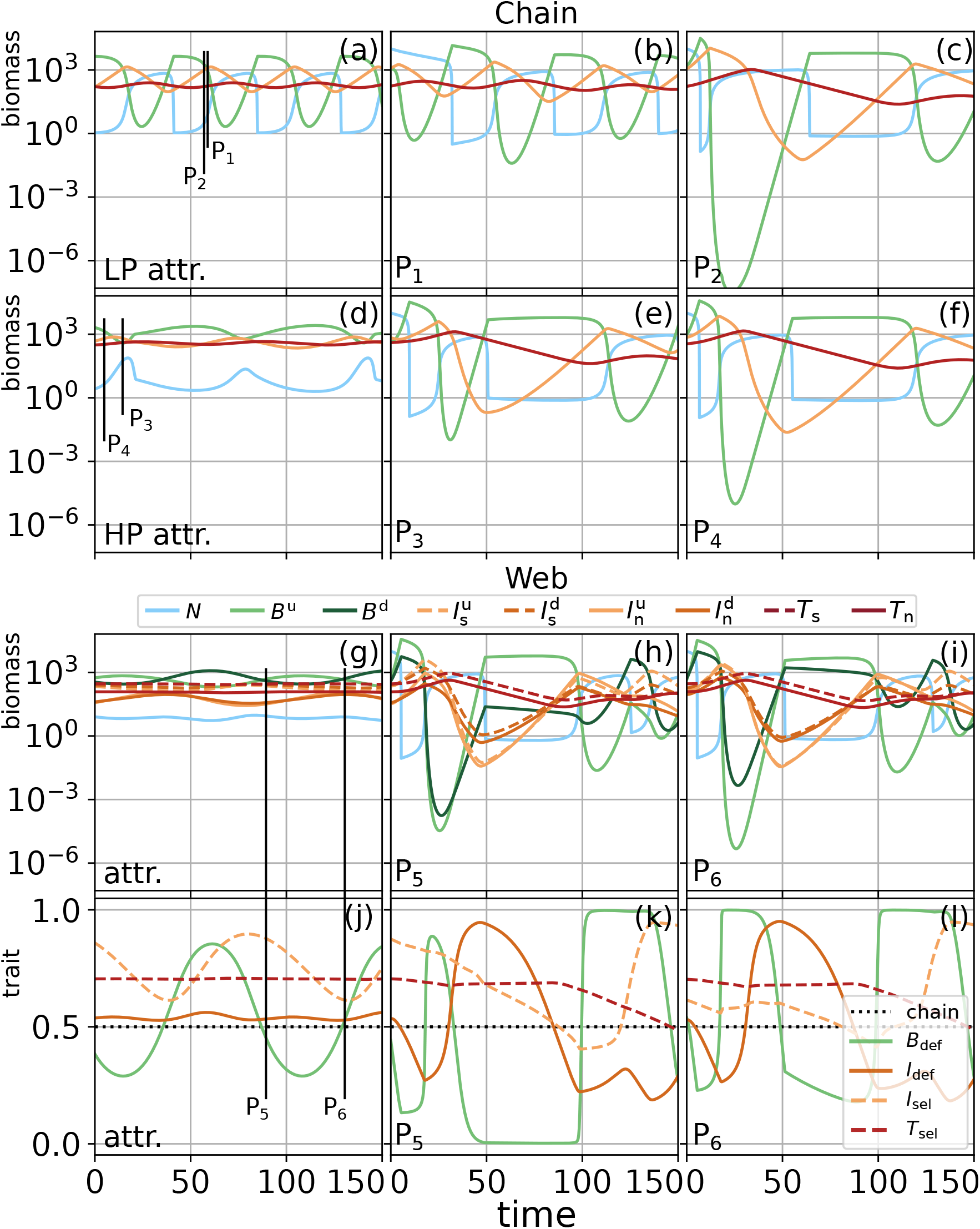
Timeseries of the dynamics of the chain and food web for *h* = 1.10, showing first the dynamics on the attractor, i.e., prior to the perturbation (chain LP: panel (a), HP: panel (d), web: panel (g)), and in the middle and right columns the system’s behavior after a perturbation of size 10^4^ *µgN / L* at different perturbation moments, here indicated by the points *P*_1_-*P*_6_ (cf. Fig. 5). The locations of these points in time on the attractor are indicated by the vertical black lines in the leftmost column. The bottom row (j–l) shows the temporal development of the trait value for the biomass dynamics shown in the panel above (basal and intermediate defense *B*_def_ & *I*_def_, and intermediate and top selectivity *I*_sel_ & *T*_sel_, cf. Fig. 1). The basal defense level *B*_def_ is the proportion of the defended basal species *B*^d^ (cf. Fig. 1) of the total amount of basal biomass. The other traits are calculated equivalently. Notably, while *P*_3_ and *P*_4_ are on the HP attractor before the perturbation, the extremely high inflow of nutrients pushes the system into the LP state, where it remains. Because the *LP* state is not attractive when *h* = 1.1 in the food web, the trajectories of *P*_5_ and *P*_6_ must eventually return to the *HP* state (cf. Fig. A3.4, Appendix).

This pattern describes what is observed generally in both the chain and the web after a nutrient pulse perturbation. A portion of the supplementary nutrients are quickly taken up by the basal trophic level, which subsequently causes a biomass peak in the intermediate trophic level. The higher this peak is, the more affected the basal level will be by the surplus grazing of the intermediate level.

A general negative correlation emerges between the maximal intermediate biomass (*I*_max_) and the minimal basal biomass (*B*_min_) shortly after the perturbation: the higher the intermediate species grow after the pulse, the more severely they deplete the basal species. Notably, our results show that a given *I*_max_ generally leads to a *B*_min_ that is approximately one order of magnitude higher in the web, as compared to the chain, and that this effect is not simply due to the different growth and grazing rates, but rather due to the increased functional diversity in the web (cf. Appendix A2 for a detailed explanation). The increase in the biomass of the intermediate trophic level leads subsequently to an increase in the biomass of the top trophic level, which in turn, leads to the intermediate level being under strict top-down control and thus unable to exploit the high basal biomass following the nutrient pulse (cf. Appendix A2, Ceulemans et al. (2019) and Ceulemans, Guill, and Gaedke (2020)). Thus, the web exhibits stronger top-down regulatory processes leading to a higher resistance to nutrient pulses.

Investigating not only the biomass but also the trait dynamics after the perturbation in the web clearly shows how a diverse food web may be able to buffer the nutrient pulse (Fig. 6j–l).Right after the perturbation, the high concentration of available nutrients causes the basal biomass to increase, with the undefended species increasing faster due to its higher growth rate. If the selective intermediate species are sufficiently high in biomass, they are able to graze down the undefended basal species to very low densities, potentially causing the defended basal species to outweigh the undefended species by several orders of magnitude (cf. Fig. 6). Importantly, the defended species is not grazed down to such low levels, preventing the potentially very strong reduction in total basal biomass observed in the chain.

Additionally, a clear hierarchy in when the trophic levels are affected is observed (Fig. 6j–l). In particular, the trait composition of the top trophic level is only substantially altered long after the perturbation. Such mechanisms can only take place in a functionally diverse community: in the linear chain, there is no potential for the mean trait value to adapt to a perturbation, and thus, no capacity for buffering.

### 3.2 Resilience

Quantifying the resilience of a food web requires determining whether the post-perturbation and pre-perturbation state differ from each other. These can differ either when the perturbation results in extinction of at least one population, or when the food web relaxes to a different attractor—on which all species coexist—in a multistable system.

Following a sufficiently large nutrient pulse, the basal trophic level biomass crosses the numerical extinction threshold of 10 ^−9^ *µgC* / *L* first, leading to additional extinctions on the *I* and *T* level (Fig. 4). When *h* = 1.1 or 1.15, a significant proportion of extinctions only happens for unrealistically large perturbation sizes outside of the range we considered. However, when *h* = 1.05, this becomes a likely occurrence starting from perturbation sizes of approximately 3. 10^3^ *µgN L* in the chain, as compared to 7. 10^3^ *µgN / L* in the web (Fig. 4). Importantly, extinctions of individual populations may already happen for smaller perturbation sizes in the web (≈ 3. 10^3^ *µgN / L*), and this leads to extinctions on higher trophic levels as well. However, the remaining population on the basal level can still support at least a part of the web. In contrast, the extinction of the basal population in the chain invariably leads to the complete disappearance of all the upper trophic levels as well. Thus, the web exhibits a higher resilience than the chain.

The above results do not account for potential differences in resilience between different attractors of the system when it is bistable (see Table 1). Previous investigation into this model showed that the basins of attraction of the high production (*HP*) and the low production (*LP*) attractor are significantly influenced by the functional diversity present: the *HP* attractor is strongly promoted as diversity increases, through efficient nutrient exploitation facilitated through compensatory dynamical patterns (Ceulemans et al., 2019).

When comparing the resilience of the high production (*HP*) and low production (*LP*) attractor to a perturbation separately, we find that the *HP* state is more vulnerable in both the chain and the web (Fig. 7). In the chain, with *h* = 1.1, perturbation sizes of maximally ≈ 1000 *µgN / L*, but even as small as ≈ 100 *µgN / L* on the *HP* state can move the system outside its basin of attraction, such that it relaxes to the *LP* state. In contrast, more points on the *LP* attractor exhibit resilience: even after perturbations of size 10, 000 *µgN / L* anywhere on the attractor, the system still returns to it (recall that *N*_0_ ≈ 1000 *µgN / L* cf Appendix A1.1).

**Figure 7.**
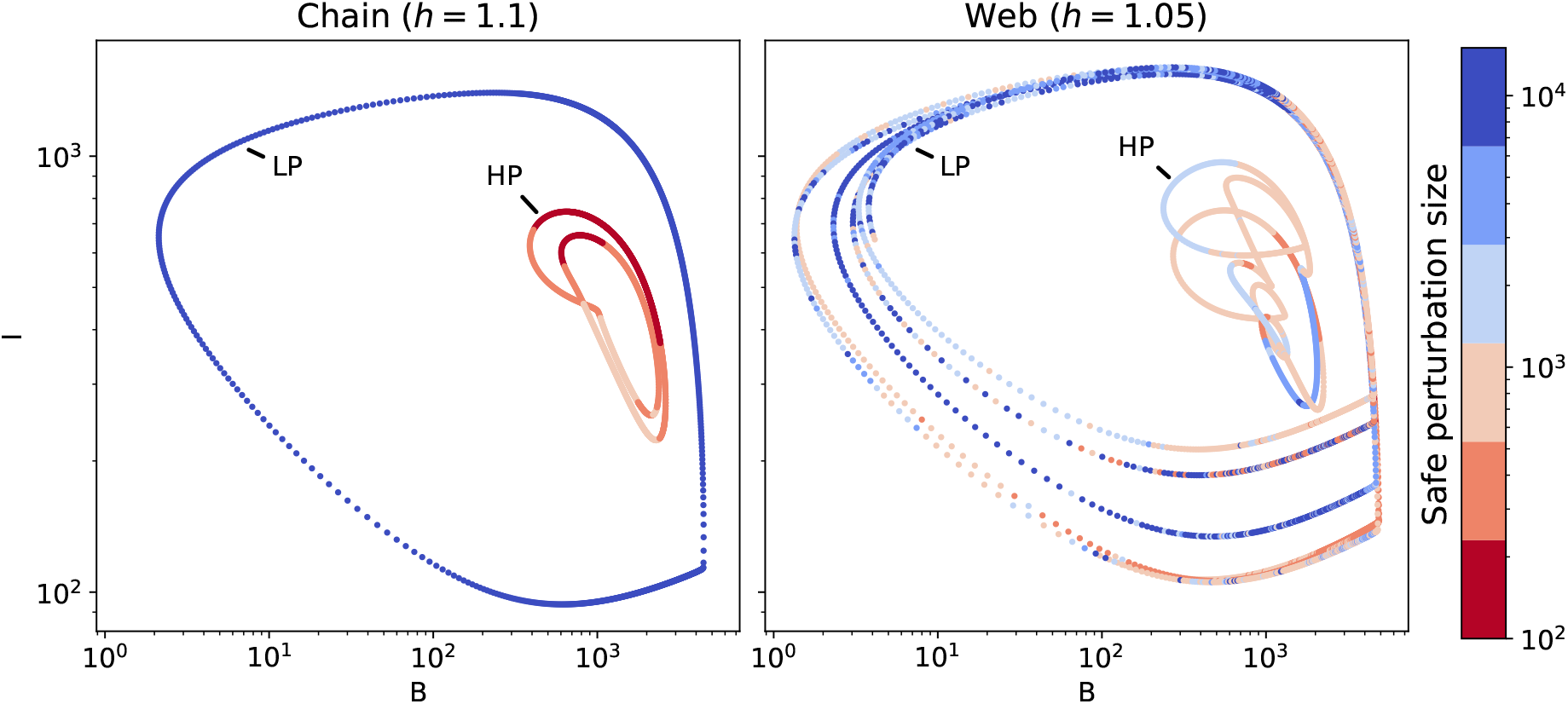
Total basal (*B*) and intermediate (*I*) biomass on the low production (*LP*) and high production (*HP*) attractors, for both the chain (left) and web (right) when they exhibit bistability. This happens when the Hill exponent *h* = 1.1 in the chain, and *h* = 1.05 in the web. The color indicates the maximum perturbation size for which the system, when perturbed at this point in the attractor, still returns to its original state. When the perturbation is larger than this “safe” perturbation size, the system either relaxes to the other attractor, or to another non-coexistence attractor. On the chain (left), the *LP* state is very resilient to perturbations, because even for perturbations of size 10^4^ *µgN / L*, the system returns to this state, independently of where on the attractor it is applied (recall that *N*_0_ ≈ 1000 *µgN* /*L*). Conversely, the *HP* state is very vulnerable: a perturbation size of ≈ 200 *µgN* /*L* frequently moves the system outside the *HP* ‘s basin of attraction, and the maximum safe perturbation is ≈ 1000 *µgN / L*. For the web, the same pattern is observed: less points on the *HP* state are resilient than those on the *LP* state. However, points on the *HP* state for the web are much more resilient those for the chain, despite its lower Hill exponent of *h* = 1.05.

For the web, with *h* = 1.05, the situation is qualitatively similar, but there are some important differences. The *LP* state still shows resilience to perturbations of ≈ 10, 000 *µgN / L*, but not over its full length. Recall that, when *h* = 1.05, the likelihood of extinctions becomes non-negligible for perturbation sizes from ≈ 3000 *µgN / L* or higher (cf. Fig. 4). Furthermore, there are some regions where perturbations of ≈ 200 *µgN / L* cause a transition from the *LP* to the *HP* state. The *HP* attractor is resilient to perturbation sizes of ≈ 5000 *µgN* /*L* for some areas, in contrast to the chain. Points *P*_3_ and *P*_4_ (Fig. 6d–f) illustrate how, when a perturbation is applied in the *HP* state, the system’s dynamics change for the *LP* state, and consequently, return to the *LP* state instead of the *HP* state. Nonetheless, our results show that functional diversity balances the transition probabilities between these two states, thereby greatly increasing the resilience of the *HP* state.

### 3.3 Elasticity

To quantify the elasticity in our system, we estimated the median return time as a function of the perturbation size (Fig. 8). This is defined as the time required for the system to stay in the attractor’s near vicinity (maximal distance to a point on the attractor must be <5 in units of total biomass) after a perturbation (cf. Fig. 2). Because this time may vary depending on where on the attractor the perturbation is applied, the median return time, as well as the lower and upper quantiles of 100 evenly spaced points on the attractor are displayed. The perturbation size is only increased up to 100 *µgN / L* (≈ 0.1*N*_0_) to prevent the influence of trajectories not returning to their original attractor.

**Figure 8.**
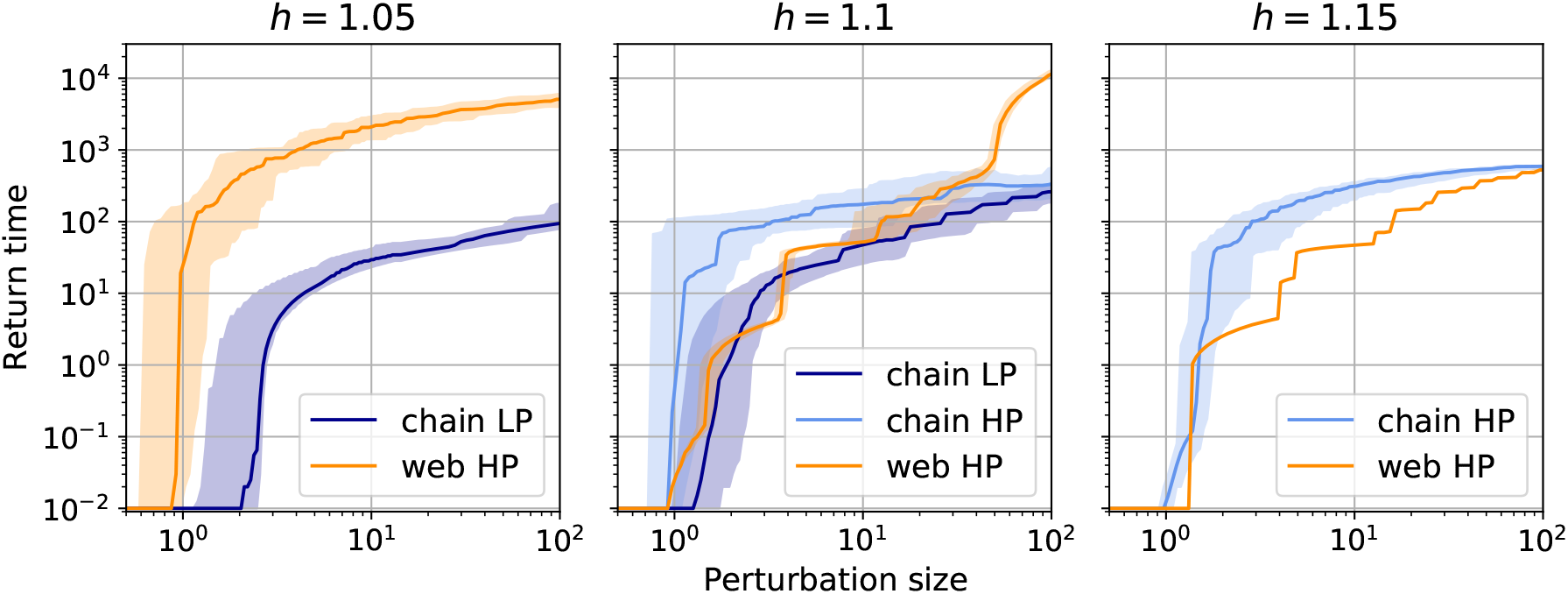
Return time as a function of the perturbation size for the different attractors in our system, for the three different Hill exponents (*h*) we investigated. Due to its complex structure, the return time for the *LP* state in the web when *h* = 1.05 (cf. Table 1) could not be accurately calculated. In all other cases, the median and lower and upper quantiles of the return times for 100 evenly spaced points on the attractor are shown for each perturbation size where applicable (when *h* 1.15, the web relaxes to a fixed point, cf. Table 1). The minimum return time (10^−2^) is determined by the time step at which the timeseries were sampled. A lower return time means that the system is faster to return to its original attractor, implying a higher elasticity. While a straightforward comparison of the elasticity of a chain versus a web proves difficult, our results show that the *LP* state in the chain tends to have a higher elasticity to nutrient pulse perturbations than the *HP* state. The elasticity of the web is higher for small perturbations than that of the equivalent state in the chain, when both exist.

Due to the significant structural differences between a chain and a web (Fig. 1), a straightforward comparison between the elasticity of these two systems is difficult. An additional complicating factor is the different number and/or type of attractors for the chain and web at a given Hill exponent. Despite these complicating factors, our results show that the return time of the *LP* attractor tends to be lower than that of the *HP* attractor. This means that the *LP* attractor tends to be more elastic than the *HP* attractor.

With this effect of the attractor shape in mind, our results show that the return time of the web tends to be shorter than that of the chain—which means the elasticity of the web is higher than that of the chain. For *h* = 1.05, the chain is on the *LP* state, while for the web, the return time could only be accurately calculated for the *HP* state (thus complicating meaningful comparison between the return time of the chain and the web). When *h* = 1.10, the web is on the *HP* state, and its return time is approximately an order of magnitude lower than the chain on the *HP* state when the perturbation is low. For higher perturbation sizes, the increased complexity of the web causes the return time to increase faster than it does for the chain. For *h* = 1.15, both web and chain are on the *HP* state, and again the return time of the web is roughly an order of magnitude lower than that of the chain. In line with expectations, the return time decreases with increasing *h* for the web, whereas this effect is less pronounced for the chain.

## 4 Discussion

We compared the consequences of a perturbation by a nutrient pulse for the dynamics of a non-diverse and a diverse tritrophic food web. The non-diverse food web is a non-adaptive food chain with three species (“chain”), whereas the diverse food web (“web”) has three adaptive trophic levels and nine species (Fig. 1). In the web, prey species can be defended or undefended against predation, and the consumer species can be selective or non-selective feeders. Changes to their fitness are balanced by two trade-offs: defended species grow slower, and non-selective feeders exploit low resource densities less efficiently.

We show that, in accordance with our first hypothesis, the higher the pulse perturbation is—and so the richer the environment becomes—the stronger the effects on the food web dynamics are (Figs. 3, 4 and 8). Consequently, temporarily increasing the available nutrients affects the trophic levels differently in our study, which conforms to the paradox of enrichment representing a press perturbation (Rosenzweig, 1971; Abrams and Roth, 1994) and widely verified by further theory (Mougi and Nishimura, 2008; Rall, Guill, and Brose, 2008), and in experimental aquatic (Persson et al., 2001) and terrestrial (Meyer et al., 2012) systems.

Additionally, as stated in our second hypothesis, the system response varies considerably depending on the moment of perturbation (i.e., the point on the attractor). In particular, we can distinguish two neighboring zones on the attractor which are either the most or the least affected (Figs. 3 and 6). This difference arises from the intermediate species ability to keep the basal level under sufficient top-down control. Under weak top-down control, the basal species are immediately able to grow to very high densities after the nutrient pulse. This basal biomass peak is followed by a peak in intermediate species biomass, which in turn leads to a depleted basal biomass level. On the other hand, if the top-down control is strong, the basal species are unable to efficiently exploit the extra nutrients, such that most of the nutrient pulse simply washes out of the system. This corresponds to previous theoretical (Rall, Guill, and Brose, 2008) and experimental findings (Weithoff, Lorke, and Walz, 2000) in a bitrophic food web, where the authors underline the importance of the top-down pressure to explain the response of systems against a nutrient perturbation. Our tritrophic study also highlights the importance of top-down processes to dampen the effects of a nutrient pulse, and reveals that the top level may strongly affect the response of the food web as a whole (see Appendix A2), because of its decisive influence on the biomasses of the two lower trophic levels (Wollrab, Diehl, and De Roos, 2012; Ceulemans, Guill, and Gaedke, 2020).

By examining different values of the Hill exponent in the basal-intermediate and intermediate-top interactions, for a chain and a web, we uncover how the type and shape of the attractor affects the system response (Fig. 3, and see Table 1). Previously, (Rall, Guill, and Brose, 2008) showed that after a nutrient increase in a simple consumer-resource system, a higher Hill exponent stabilized the population dynamics. We confirm this pattern for tritrophic systems and show additional effects of the shape and type of attractor present in the complete state space (Fig. 3).

### Resistance generally increases with functional diversity

Considering each trophic level as a whole, the web is generally more resistant, as stated in our third hypothesis, since the biomasses do not reach as low values as in the chain (Figs. 4 and 5). However, at the population level, the undefended basal species (*B*^*u*^) may be more affected in the web than the only basal species in the chain for instance (Figs. 4 and 6f, i). In other words, the system’s resistance varies with the organization level studied, i.e., the trophic or population level.

The above analysis rests on the general negative relationship between the minimal biomass of the basal trophic level (*B*_min_) and the maximal biomass of the intermediate trophic level (*I*_max_) after a nutrient pulse perturbation (Figs. 3 and see Appendix A2). It reveals that a higher *I*_max_ leads to a lower *B*_min_ due to the increased grazing of the basal level. Importantly, the web maintains a considerably higher *B*_min_ at a given *I*_max_, compared to the chain (see Appendix A2a), even when accounting for the effects of different parametrizations (see Appendix A2b, c).

Under the extremely nutrient-rich conditions immediately following the perturbation, *B*^*u*^ is at a competitive advantage due to its higher growth rate, which explains its dominance during the basal biomass peak over the entire trophic level. As a consequence, the main consumers of *B*^*u*^ strongly increase, which in turn leads to *B*^*u*^ being grazed down to very low biomasses. On the other hand, because of its slower growth rate and competitiveness the basal defended species, *B*^*d*^, only experiences limited additional growth, and thus only contributes little to the growth of its intermediate consumers. In turn, *B*^*d*^ is grazed down less thanks to its lower growth rate and stabilizes the trophic level biomass (Fig. 6i, l). Therefore, we highlight how a defended (slow-growing) species in a tritrophic food web increases the resistance of the corresponding trophic level. This is in line with previous studies considering defended species in tritrophic food chains (Loeuille and Loreau, 2004) or slow-growing plant species (Oliver et al., 2015).

In this way, our model reveals the mechanism behind how species’ functional traits determine their dynamics in a food web. This leads to explicit manifestations of the insurance hypothesis (Naeem and Li, 1997), since a higher resistance is observed at the aggregated trophic level for the web than for the chain, thanks to the average response of different species.

### Nutrient pulse pushes system to low-production state affecting the system’s resilience

The resilience of a system can be affected by extinctions, or by the presence of alternative stable states in which all species coexist. Our results show that the resilience of both the web and the chain strongly depends on the moment of perturbation (Fig. 7). We find that extinction of a complete trophic level is protected by functional diversity, however, extinction of a single species is equally probable in the chain as in the web (Fig. 4). The crucial difference is that species extinction in the chain necessarily causes the secondary extinctions of all species at higher trophic levels. In contrast, after an extinction of a single species in the web, much of the trophic structure persists due to its functional redundancy (Borrvall, Ebenman, and Jonsson, 2000; Fonseca and Ganade, 2001). Generally, we show that species extinction occurs regularly in the chain and in the web for perturbation sizes over 10^3^ *µgC* / *L*. However, even for much smaller perturbation sizes, both food chain and food web may be vulnerable to a regime shift when the perturbation causes the system to relax to another coexisting attractor.

In our model, both the chain and the web exhibit bistability for a large part of the parameter space (Table 1). In these cases, the system can relax to either the low-production (*LP*) or high-production (*HP*) state, depending on the initial conditions. This implies that a sufficiently large perturbation can result in the system relaxing to the other state, which affects its resilience. Such behavior, commonly called a regime shift, is a widely observed phenomenon that can occur in many different types of ecosystems (Scheffer and Carpenter, 2003; Folke et al., 2004). Regime shifts are often the cause of major concern, because the two states may vary considerably in their ecological properties. Some examples are changes in vegetation patterns (Dublin, Sinclair, and McGlade, 1990), in particular under desertification (Bestelmeyer et al., 2015); or transitions between a clear and a turbid state in lakes (Scheffer et al., 1993; Scheffer and Jeppesen, 2007).

We previously showed that diversity loss likely causes our system to transition from the *HP* state to the ecologically undesirable *LP* state, where top level biomass is much lower, and the biomass dynamics more variable (Ceulemans et al., 2019). Here, we showed that the system is more likely to transition to the *LP* than to the *HP* attractor, when exposed to a sudden nutrient pulse (Fig. 7). This asymmetry can be explained by a closer analysis of the post-perturbation timeseries shown in Fig. 6. After a nutrient pulse, both basal species increase due to the high amount of readily available nutrients. As a response, all intermediate species increase, which causes a concurrent decrease of all basal species, and increase of all top species. In other words, immediately after a perturbation, species on the same trophic level tend to move synchronously, largely independently of their trait values. This is exactly the dynamical pattern that governs the behavior of the *LP* state. In contrast, on the *HP* state, the species tend to exhibit compensatory dynamical patterns (Fig. 6g, and (Ceulemans et al., 2019)). Therefore, a nutrient pulse causes the system to behave like the *LP* state, and a relatively small disturbance of approximately *N*_0_ (the normal inflow nutrient concentration), or less, forces the system to stay permanently in this state. Interestingly, the low resilience of the *HP* state, together with its low temporal variability of the biomass dynamics (and vice-versa for the *LP* state), means that our model explicitly shows how the different aspects of stability may co-vary with each other: being very stable in one aspect may come at the price lesser performance in another aspect (Donohue et al., 2016; Domínguez-García, Dakos, and Kéfi, 2019).

In summary, a nutrient pulse can cause a regime shift between the *HP* and *LP* attractors, mostly from the *HP* to the *LP* attractor. Combined with our knowledge of resistance, our findings highlight the key role played by functional diversity in governing the response of a food web. When functional diversity is high, the *HP* state persists. The high top biomass level then ensures adequate control on the intermediate level, in turn protecting the basal level from over-exploitation. A reduction in functional diversity can therefore abruptly affect food web resilience, as the system is more easily kicked to the *LP* attractor, where top level biomass is low.

### Elasticity depends on attractor structure, shape, and diversity

Another way to quantify a system’s response after a perturbation is by measuring its elasticity, that is, the time it takes to return to the pre-perturbation state (return time, cf. Fig. 2). In an economical context, elasticity is an important quantity, because low elasticity means that the desired functioning of an ecosystem may be interrupted for a substantial period of time before returning back to normal (Oliver et al., 2015).

Our results show that the return time increases with the size of the nutrient pulse, and, moreover, this increase can happen in discrete jumps: suddenly, the dynamics require almost a whole additional revolution in state space before being close again to the attractor (Fig. 8). We also observed that the return time depends on functional diversity and the shape of the attractor to which the perturbed system is returning. In particular, we find that the return time for the *HP* state tends to be higher than for the *LP* state, which is due to the different shapes of the two attractors. On the *LP* attractor, individual species’ amplitudes are significantly higher than on the *HP* attractor, e.g., both the nutrients and the basal trophic level routinely reach densities close to their carrying capacity. This dynamical behaviour is very similar to that of the immediate post-perturbation dynamics (cf. Fig. 6). Moreover, perturbing the *HP* state can lead to the dynamics permanently (cf. Fig. 7) or temporarily (cf. Figs. 6 and A3.4) behaving like the *LP* attractor. Notably, this can happen even when the *LP* state is not a dynamical attractor (cf. Fig. A3.4). In this case, the time spent by the transient on this ghost attractor increases with the distance to the bifurcation (Hastings et al., 2018; Morozov et al., 2020), ultimately strongly affecting the return time of the *HP* state as the Hill exponent decreases. A meaningful comparison between food webs of low and high diversity thus requires precise knowledge about the system’s attractor(s). This observation highlights the difficulties in generalizing the diversity-stability relationship: in realistic ecological systems this relationship and its corresponding mechanisms may become highly complex (McCann, 2000; Ives and Carpenter, 2007; Loreau and Mazancourt, 2013). Notwithstanding these complexities, our results suggest that the return time is lower in the web than in the chain, i.e the web is more elastic than the chain (Fig. 8). Our study thus confirms the positive relationship between elasticity and functional diversity observed for one trophic level in other studies (Smith, Vallina, and Merico, 2016; Schmitt et al., 2020) and extends it to tritrophic food webs.

### Concluding remarks

Overall, this study reveals that tritrophic systems are more strongly affected by a pulse perturbation of higher magnitude, and that the moment when the perturbation occurs determines the consequences for the following dynamics of the system. Importantly, we show how functional diversity buffers the effects of a perturbation: increased functional diversity leads to a higher resistance, resilience and elasticity, and dampens the risk of inducing a regime shift towards an ecologically less desirable stable state. Even though a nutrient pulse only directly affects the basal trophic level, we reveal how top-down regulatory processes determine the system response. We thus uncover the role of both horizontal (i.e., functional diversity within each trophic level) and vertical diversity (i.e., the number of trophic levels) in governing how food webs respond to disturbances. In this way, the potentially destructive positive feedback loop is mechanistically understood: a loss in functional diversity affects food web functioning in such a way that its resilience, resistance and elasticity become lower, making the food web even more vulnerable to future perturbations.

## Acknowledgements

We thank George Adje, Elias Ehrlich, Guntram Weithoff, Lutz Becks, and Nicolas Loeuille for giving feedback on an earlier version of the manuscript. This project was funded by the German Research Foundation (DFG) Priority Programme 1704: DynaTrait (GA 401/26-2).

