## Supplementary material for "Functional diversity alters the effects of a pulse perturbation on the dynamics of tritrophic food webs"

*: shared first authorship

University of Potsdam, Germany

### A1 Additional information about the model

| Parameter name | Parameter value | |
|---|---|---|
| Body mass ratio between adjacent TLs | $m_I/m_B = m_T/m_I = 10^3$ | |
| Allometric scaling exponent | $\lambda = -0.15$ | |
| Inflow nutrient concentration | $N_0 = 1120\mu gN/L$ | |
| Dilution rate | $\delta = 0.055$ | |
| Nutrient half-saturation const. of B | $h_N = 10\mu gN/L$ | |
| Nitrogen to carbon ratio of B | $c_N/c_C \approx 0.175$ | |
| conversion efficiency | $e = 0.33$ | |
| T max. grazing rate | $\gamma' \approx 0.38/day$ | |
| Hill exponent | $h = \{1.05, 1.10, 1.15\}$ | |
| Parameter name | chain | web |
| B max. growth rate | $r'_c \approx 0.81/day$ | $r'_u = 1/day$, $r'_d = 0.66/day$ |
| I max. grazing rate | $g'_c \approx 0.87/day$ | $g'_u \approx 1.08/day$, $g'_d \approx 0.71/day$ |
| B-I half-saturation const. | $M_c \approx 424\mu gC/L$ | $M_s = 300\mu gC/L$, $M_n = 600\mu gC/L$ |
| I-T half-saturation const. | $\mu_c \approx 424\mu gC/L$ | $\mu_s = 300\mu gC/L$, $\mu_n = 600\mu gC/L$ |

Table A1.1: Parameters values used as described in the tritrophic chemostat model developed by Ceulemans et al. (2019). The chain and the web have different parametrizations according to the trade-offs between the relevant traits. As a consequence, the $B$ and $I$ maximal growth rates, as well as the $B$–$I$ and $I$–$T$ half-saturation constant decrease with the defense for the prey and the selectivity for the predators, respectively. The trait values of the standard chain are included in the range of the values set for the web, such that $r'_u > r'_c > r'_d$, $g'_u > g'_c > g'_d$, $M_s > M_c > M_n$ and $\mu_s > \mu_c > \mu_n$. Simulations of the chain and the web are compared for three different values of the Hill exponent.

Following (Ceulemans et al., 2019), the exact values of the trait values in the chain are calculated from those in the diverse web as follows:

$$\log \theta_c = \frac{\log \theta_a + \log \theta_b}{2}, \qquad (1)$$

where $a = u$ and $b = d$ for $\theta = r'$ or $\theta = g'$; or $a = s$ and $b = n$ for $\theta = M$ or $\theta = \mu$.

### A2 The $B_{\min} - I_{\max}$ relationship

In this section, we provide more details about the relationship between the minimal basal biomass ($B_{\min}$) and the maximal intermediate biomass ($I_{\max}$) reached after the perturbation, that we describe in the main text. We verify that this negative relationship holds for different parametrizations (Hill exponent, basal growth rate and intermediate half-saturation constant). Particularly, we investigate how functional diversity influences this $B_{\min} - I_{\max}$ relationship by comparing the food web alongside differently parametrised food chains, and we also uncover the principal role of the top trophic level in governing the response to a nutrient pulse perturbation in tritrophic systems (Fig. A2.1).

The negative relationship that is observed between $B_{\min}$ and $I_{\max}$ is influenced by several food web parameters, such as the Hill exponent (Fig. A2.1a): lowering the Hill exponent increases the oscillation amplitude of the dynamics, such that food webs with a Hill exponent $h = 1.05$ at the low-production (LP) state obtain the lowest $B_{\min}$ values. We also observe that the high-production (HP) state has lower $I_{\max}$ and higher $B_{\min}$ values, as compared to the LP state (Fig. A2.1a). Because the HP state is characterized by a higher top species biomass and smaller amplitudes (Fig. 6a and d), it does not reach the extreme values of the LP state (Fig. 6c and f). Moreover, the top species keep the intermediate species under stronger top-down control, which limits their response to nutrient enrichment.

Importantly, our results show that $B_{\min}$ tends to be higher in the food web than in the food chain (Fig. A2.1a). A given $I_{\max}$ leads to a higher $B_{\min}$ in the food web (insofar as they can be compared). This suggests that a diverse food web is more resistant to a pulse perturbation than a food chain with little or no diversity.

To untangle a potential direct effect of diversity on the $B_{\min} - I_{\max}$ relationship from the effect of comparing basal and intermediate species with different growth rates and interaction parameters in the chain and web, we also compare the food web to differently parametrized food chains (Fig. A2.1b). The maximal basal growth rate in the food chain ($r'_c \approx 0.81$) lies in-between the defended basal growth rate ($r'_{\mathrm{d}} = 0.66$) and the undefended basal growth rate ($r'_{\mathrm{u}} = 1$) of the diverse food web. Additionally, the basal-intermediate half-saturation constant in the food chain ($M_c \approx 424$) also lies in-between the values of the basal-selective ($M_{\mathrm{s}} = 300$) and basal-non-selective ($M_{\mathrm{n}} = 600$) interactions of the food web. Evidently, changing these parameters in the food chain also modifies the resulting $B_{\min} - I_{\max}$ relationship, either through direct (such as grazing suppression at low prey densities when $M$ is high), or indirect effects (such as differing top biomasses when the basal growth rate $r$ is changed, Fig. A2.1c).

Importantly, in the food web, the growth rates of the undefended and defended $I$ species differ from the growth rate of I in the food chain. However, in neither a chain where all growth rates are scaled to reflect that of the defended species, nor of the undefended species, is it possible for the top trophic level to survive. We therefore compare the food web to a chain where only the basal growth rate is altered. In the food chain with $r_B$ set to the high value of the undefended species in the web ($B^{\mathsf{u}}$), $B_{\min}$ remains approximately one order of magnitude below $B_{\min}$ of the food web, whereas decreasing $r_B$ to the value for the defended species decreases ($B^{\mathsf{d}}$), $B_{\min}$ (Fig. A2.1b). These changes are due to strong differences in the mean biomass on the top level in the alternative food chains: when $r_B$ is high, the increased basal productivity translates to an increased biomass in the top level, and vice-versa (cf. Figs. A2.1c and A2.2). When the top biomass is higher, stronger grazing pressure on the intermediate level prevents excessive grazing of $B$. On the other hand, for low top biomass the intermediate level can remain at high biomass for an extended period of time, until they find no more food.

Similarly, increasing the $B - I$ half saturation constant in the chain to the value of the non-selective consumers in the web ($M = 600$) increases $B_{\min}$; and decreasing $M$ to the value of the selective consumers in the web (300) correspondingly decreases $B_{\min}$. These changes are caused by the grazing suppression at low $B$ densities, when $M$ is increased. However, the changed mean top biomass caused by the altered basal productivity dominates the changes in $B_{\min}$ when varying $M$ (cf. Figs. A2.1c and A2.2).

In the food web, $B^u$ is generally grazed to lower densities than $B^d$ (Fig. 6h, i). This means that the $B^d - I_n$ interaction (low $r$, high $M$) is principally responsible for the value of $B_{\min}$ (Fig. 1). Remarkably, in a chain parametrized to have this interaction, $B_{\min}$ is still approximately one order of magnitude below that of the food web.

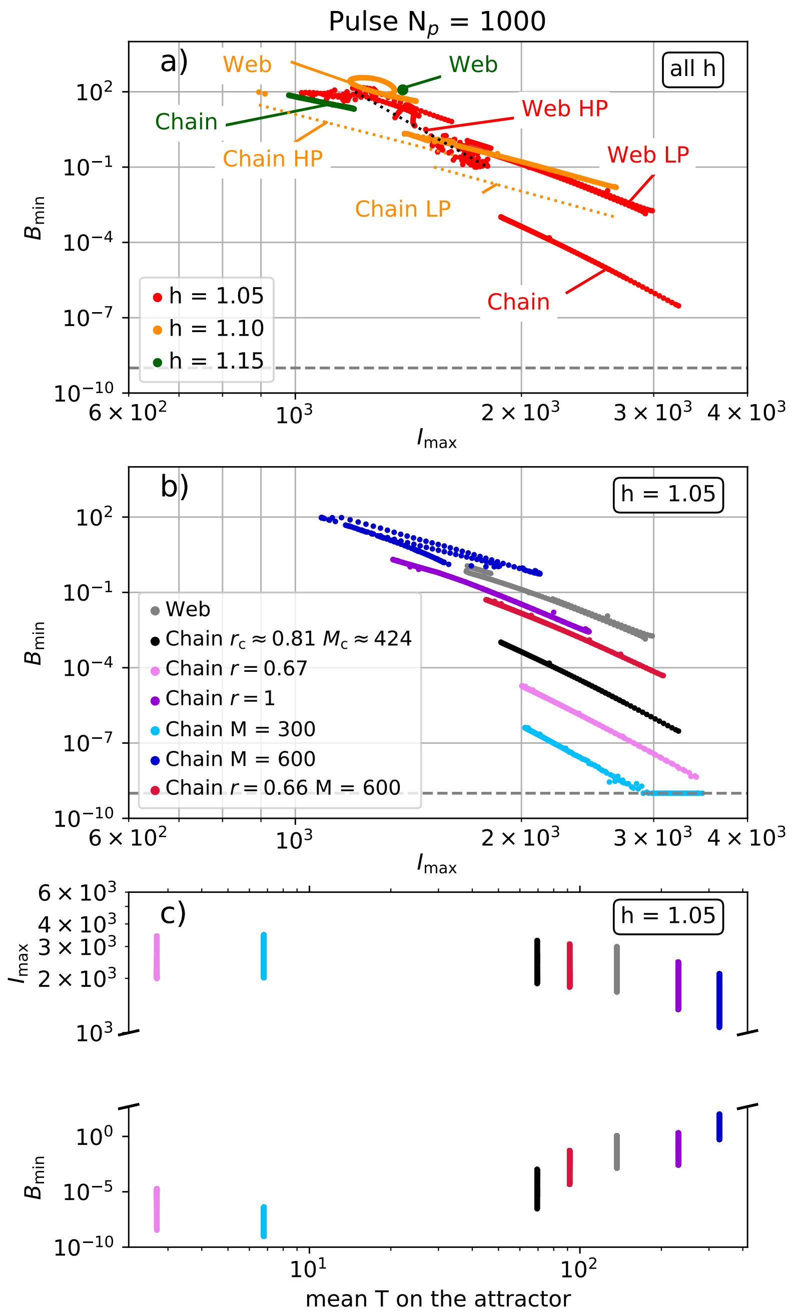


Figure A2.1: Relationship between the minimal biomass reached by the basal level ($B_{\min}$), and the maximal biomass reached by the intermediate level ($I_{\max}$), after a nutrient pulse perturbation $N_{\mathrm{p}} = 1000$, depending on where on the attractor the perturbation is applied. The collections of points in the graph are grouped per attractor (LP: low-production state, HP: high-production state, cf. Table 1, main text). A general negative correlation between $B_{\min}$ and $I_{\max}$ can be observed: the higher the intermediate level is able to grow after the perturbation, the lower it will graze down the basal level. This relationship becomes more pronounced for lower Hill exponents, as the amplitude of the dynamics increases (panel a). Importantly, when comparing the food web to the food chain while keeping the Hill exponent constant, the same $I_{\max}$ leads to a higher $B_{\min}$ in the food web. This ratio is influenced by the growth rate of the basal species and the half-saturation constant of the basal-intermediate interaction (panel b) as well as the biomass of the top trophic level on the attractor before the perturbation (panel c). Here, the growth rate of the basal species only is set to $r_{\mathrm{B}}$ (standard $r_{\mathrm{B}} \approx 0.81$), and/or the $B - I$ half-saturation constant is set by $M$ (standard $M \approx 424$).

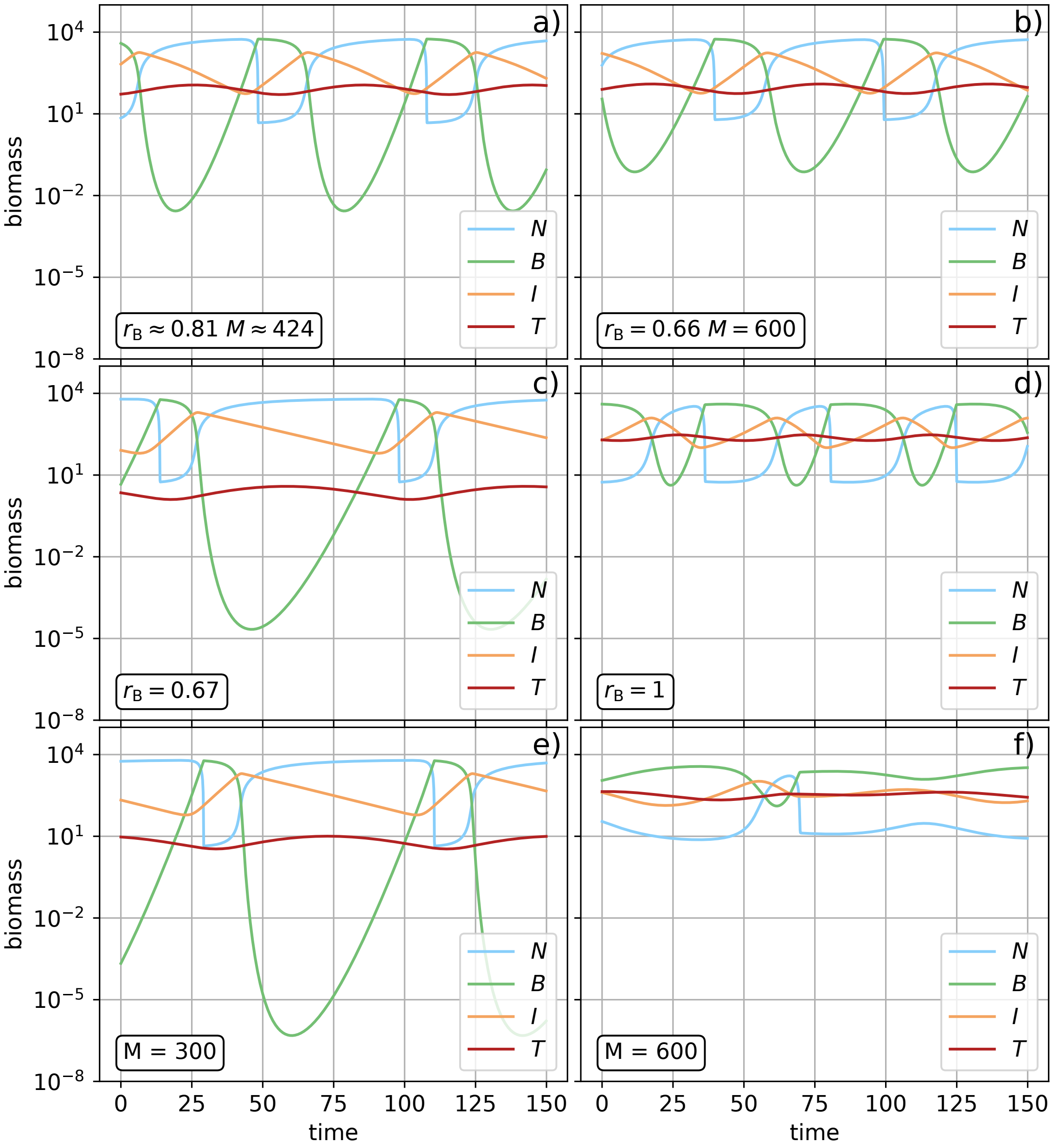


Figure A2.2: Timeseries of the dynamics on the attractor for the differently parametrized food chains in Fig. A2.1. Panel a shows the standard parametrization of the chain where $r_b$ and $M$ are logarithmic in-between the defended and undefended growth rate, and selective and non-selective half-saturation constant, respectively. Panels c and d show how the dynamics are affected by setting $r_B$ to that of the defended basal species (c), and undefended basal species (d). The change in basal biomass production directly translates to a change in biomass on the top level, which in turn changes top-down control on the intermediate level. Panels e and f show how changes to the basal-intermediate half-saturation constant $M$ has a similar effect on the top level, and thus, indirectly also on the basal level.

#### A3 Additional figures

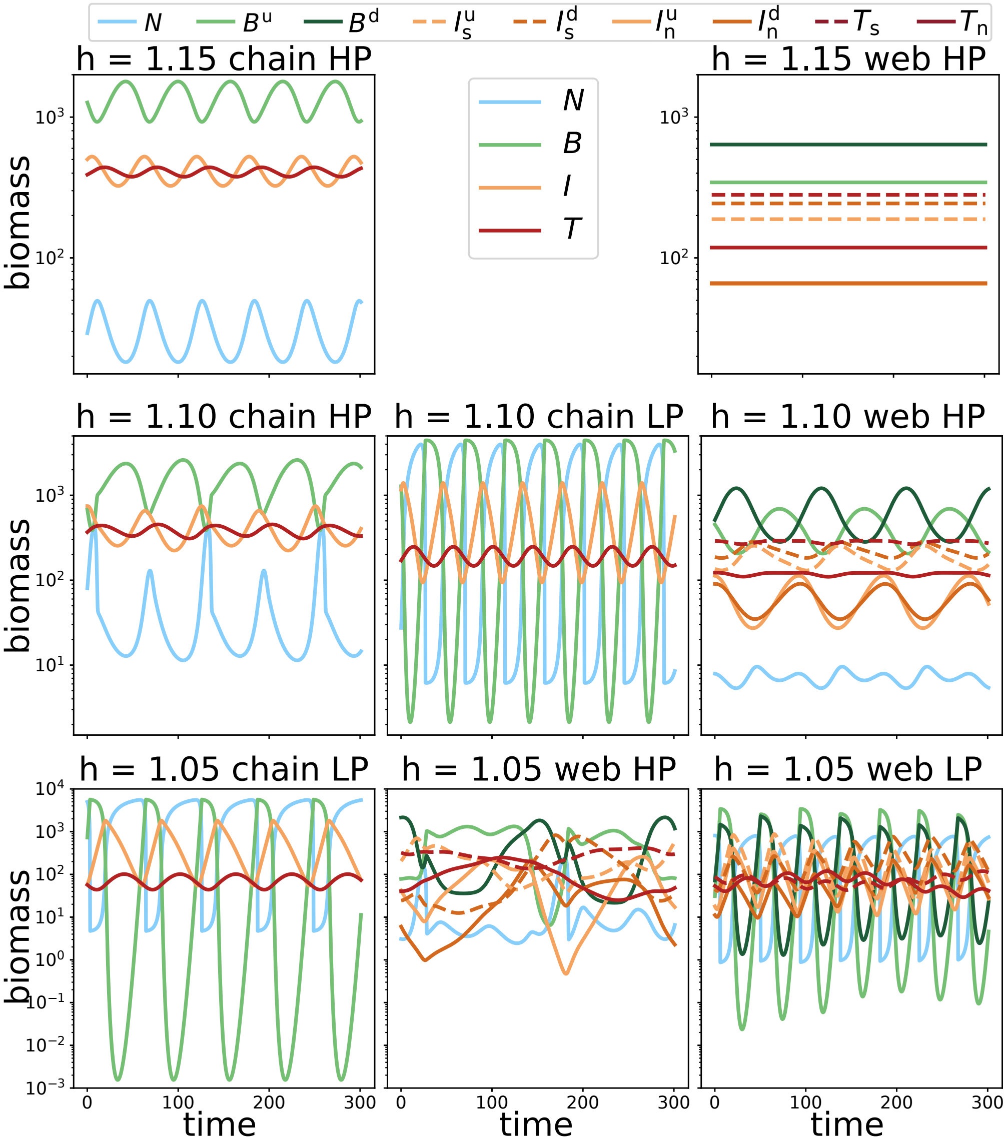


Figure A3.1: Short overview of the dynamics on each of the different attractors. The type and shape of the attractors depend on the Hill exponent h (cf. Table 1, main text). Note the different biomass scales per row. The top legend only applies to panels in which the dynamics of the food web are shown. Notice how the high production ($HP$) attractor is characterized by a low mean free nutrient concentration, high top biomass, and generally low temporal variability, in contrast to the low-production ($LP$) attractor.

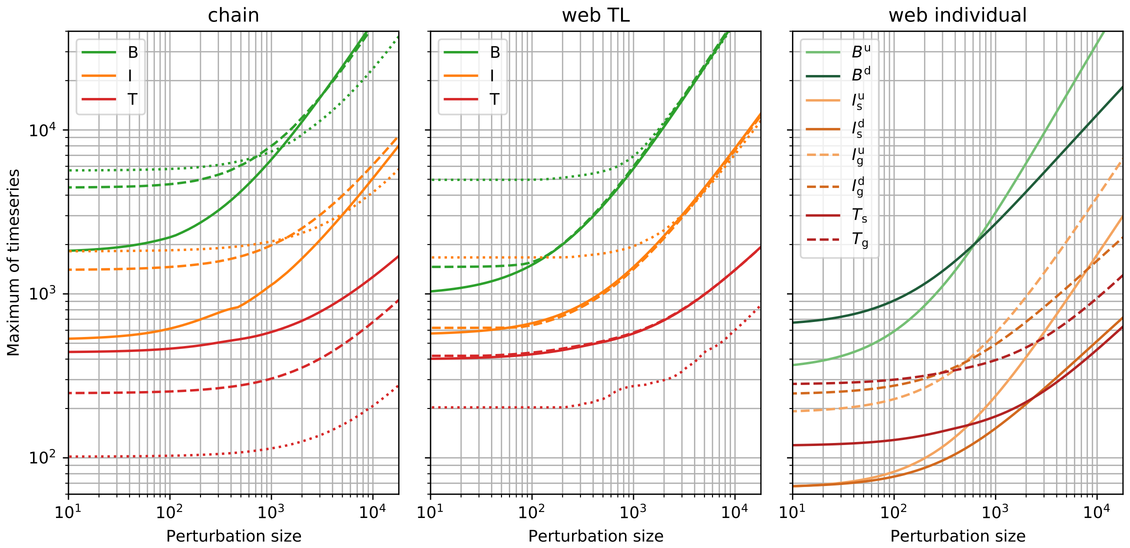


Figure A3.2: Biomass maxima reached by the timeseries after the perturbation, for the chain (left), the total biomass per trophic level in the food web (middle), and the individual populations in the food web (right), as a function of the perturbation size. Each line corresponds to the median of 1000 different randomly sampled initial conditions that lead to coexistence of all species, with the shaded area showing the upper and lower quantiles. These initial conditions were first allowed to relax to the attractor for $3 \cdot 10^4$ time units before the perturbation was applied. For the chain and trophic level biomass in the food web, the solid lines show the maxima for Hill exponents $h = 1.15$, dashed for $h = 1.10$, and dotted for $h = 1.05$. The individual populations maxima for the food web are only shown in the case of $h = 1.05$.

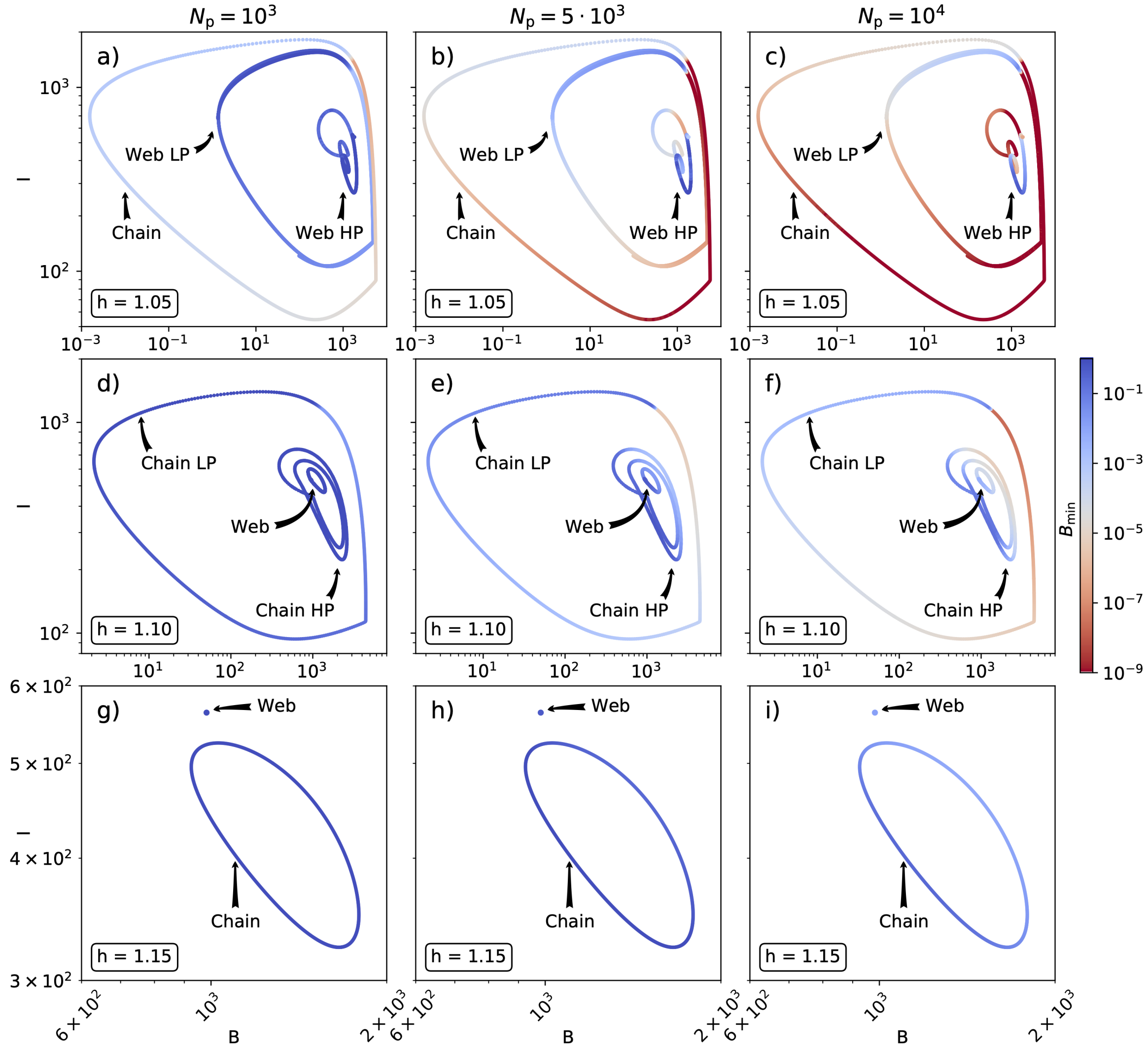


Figure A3.3: Biomass minima reached by the basal species after a perturbation, $B_{\min}$ depending on its location on the attractor, for different perturbation sizes. The general pattern that the web and $LP$ state resistance tend to be higher than the chain and $HP$ state, respectively, holds for different perturbation sizes and different Hill exponents.

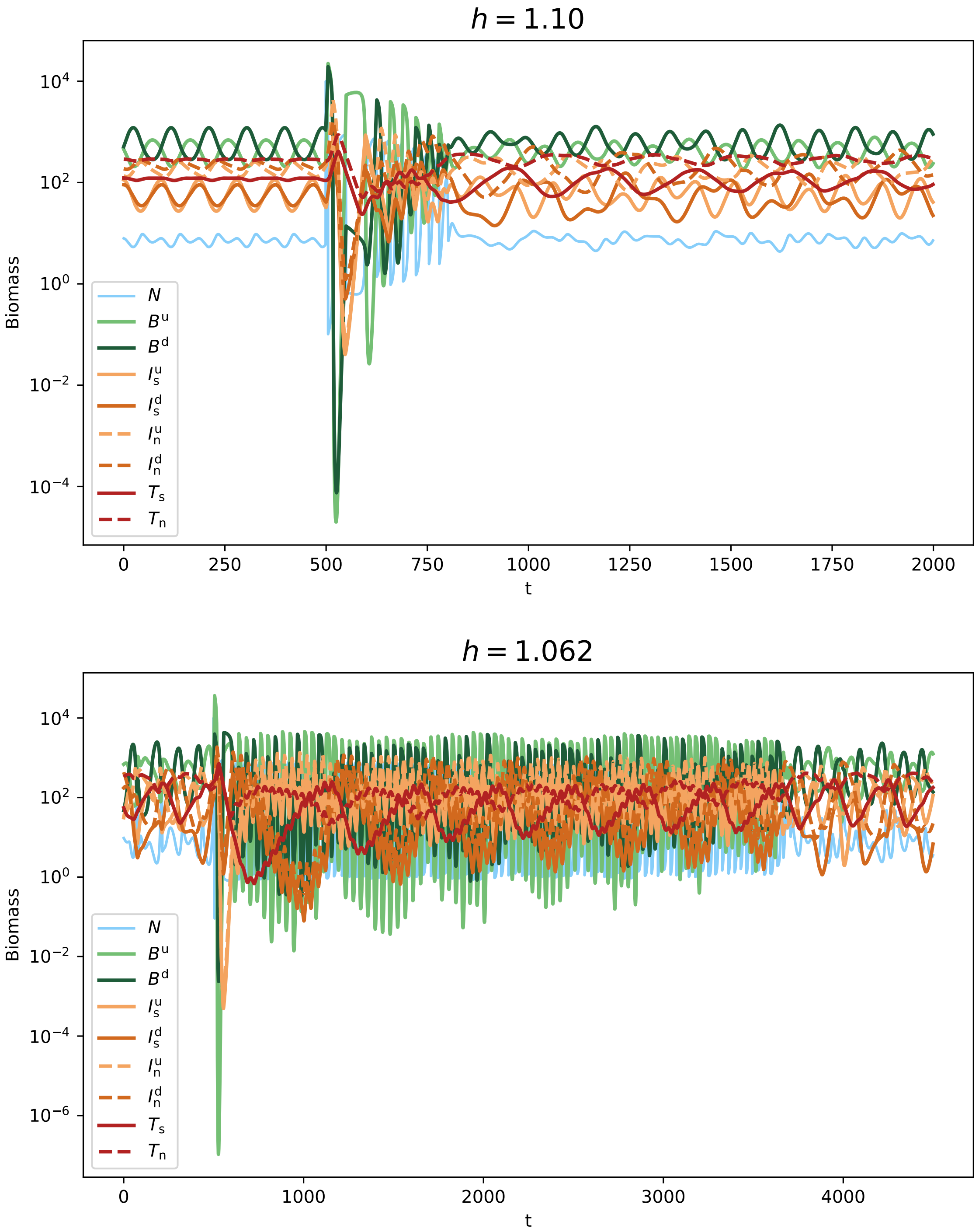


Figure A3.4: The ghost attractor phenomenon in the post-perturbation transient. When $h = 1.10$, the only stable state in the food web is the $HP$ state (cf. Fig. A3.1, and Table 1, main text). However, when perturbed by a nutrient pulse (in this case $N_P = 10^4$ at $t = 500$), the dynamics temporarily behave like the $LP$ state (see also Fig. 6, main text). After approx. 250 time units, the dynamics shift again to that of the $HP$ state, but require a long time to settle down to regular oscillations. The $LP$ state becomes unstable for approximately $h > 1.06$ (Ceulemans et al., 2019). Close to this threshold, the dynamics may be on the ghost state for much longer after the same perturbation (note the different time scales in the plots).
